# Equity in Action: A Four-Year journey towards Gender Parity and Racial Diversity in Biochemistry Hiring

**DOI:** 10.1101/2024.10.10.617211

**Authors:** Sherri L. Christian, Valerie Booth, Scott Harding, Amy M. Todd, Mark D. Berry

## Abstract

Recruitment of faculty members in academic departments shapes the department for decades in both research and teaching arenas. Having a diverse department is beneficial for undergraduate and graduate students as representation of underrepresented minority groups in the professoriate can inspire a greater diversity of students to pursue higher levels of education or research-focused careers. Increased diversity benefits research directly as diverse teams have been shown to have better ideas and outcomes. In 2020, our department had lower gender diversity than would be expected based on the pool of PhD students and post-doctoral fellows in Canada. Therefore, we altered our hiring process, primarily by redacting applications, for recruitment into entry-level tenure-track faculty positions. With this change in process, female hires increased from 17% in the previous ten years (5 hires) to 80% in the subsequent four years (5 hires) with no substantial change in hiring of racially diverse individuals (50% to 40%). Overall, combined with retirements, the percentage of female faculty in the department went from 25% to 50% and the percentage of racialized faculty went from 38% to 44%. The new hires have met or exceeded expectations of success with respect to grant funding and are on track to meet or exceed expectations for other metrics of success. Thus, our intervention was very successful in increasing the diversity of our department within a short timeframe. We believe that our experience could provide other departments with a template for making substantive change, even in the absence of internal expertise in the area.

**Graphical Abstract:** 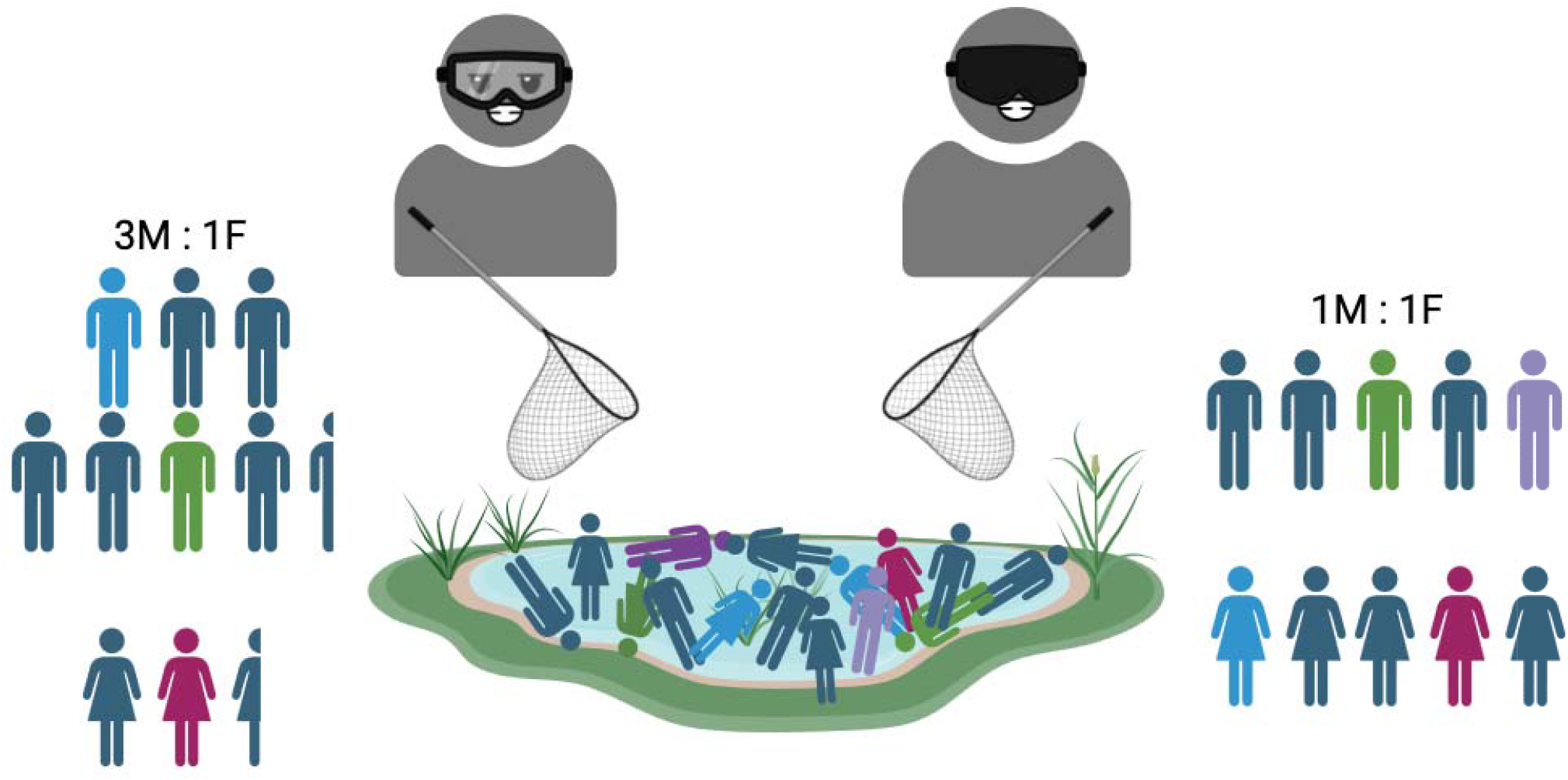

## Introduction

In academia, the process of faculty recruitment plays a crucial role in shaping the future of educational institutions. Faculty hiring processes, however, have often been criticized for being susceptible to implicit biases ^1^, which can inadvertently influence hiring decisions. Biases can stem from various sources, including the candidate’s name, sex, gender, race, ethnicity, age, accommodation requirements, and the prestige of previously attended institutions. Multiple studies have clearly shown that identical Curriculum Vitae (CV) with different names are evaluated differently with implied race, ethnicity, and gender contributing substantially to these dissimilar evaluations ^2–6^. It has been modeled that even a 3% evaluation bias at each career stage can result in a significant (∼40%) reduction in diversity over time ^1,7^. Overall, these implicit biases can overshadow the true merit and potential of applicants, leading to less diverse and less inclusive academic environments. This is to the detriment of scientific progress as diverse teams have been shown to have better ideas and outcomes than teams formed of similar individuals ^8–13^.

The issue of implicit bias in faculty hiring came to the forefront when we recognized that our department had substantially more male faculty than female faculty at the end of 2020 (Figure 1). This became particularly apparent after a period of approximately 10 years where only one out of five (20%) recruitments were female. This is low compared to the fraction of female scientists expected to be in the pool of candidates, which can be estimated from the applications for postdoctoral fellowships through the Natural Sciences and Engineering Research Council of Canada (NSERC) in the areas relevant to our department. We believe that the comparison to post-doctoral fellowship applications is the most relevant because these individuals have indicated a clear interest in continuing in academic research at a University, College, or Research Centre. In 2021, female scientists made up 53%, 36%, and 31% of applicants to Cellular and Molecular Biology, Plant and Animal Biology, and Chemistry evaluation groups, respectively ^14^.

**Figure 1.**
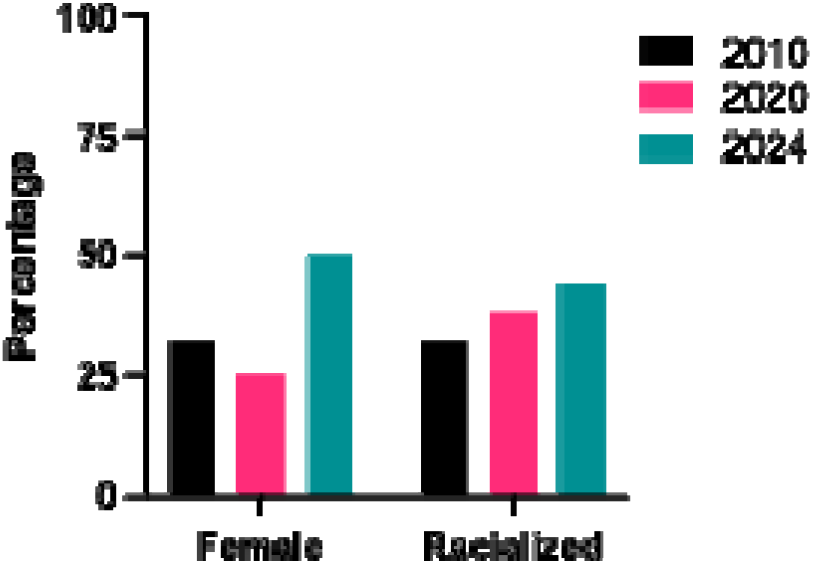
Makeup of the Department in terms of female and racialized individuals. The percentages of female and racialized faculty are shown in 2010, 2020 and 2024, which bookend two periods of analysis.

The motivation to actively change our processes came from data suggesting faculty-hiring patterns are not parallel to the availability of female candidates. The proportion of recently qualified scientists who are female has been near parity in fields related to Biochemistry since 2004 (Figure S1) ^15^. In 2020 (the same year that PDFs would enter the 2021 competition), 46.4% of people graduating with PhDs in “Physical and life sciences and technologies” and 62.9% in “Health and related fields” in Canada were women ^15^. This was similar in 2010, where 42.9% and 60.6% of these groups were women, respectively. Importantly, women have represented a large fraction of young scientists in our field for several decades, but are not being hired as faculty in corresponding numbers. Waiting for faculty-hiring statistics to correct on their own has not worked; therefore, changes need to be made to the hiring processes themselves.

We were unable to find similar statistics for racialized groups in science as these numbers have historically been poorly tracked in Canada (https://www.macleans.ca/opinion/canadian-universities-must-collect-race-based-data/), but we did find that, according to Statistics Canada, 9.2% of Canadians in a visible minority in 2021 and 7.5% of Canadians in a visible minority in 2011 held a “University certificate, diploma or degree above bachelor level” ^16^. Therefore, while women were underrepresented in our faculty complement, racialized individuals seemed to be better represented.

Therefore, as a department, we decided to initiate changes to our hiring processes to decrease implicit bias in order to increase the diversity of the faculty complement with an emphasis on sex-diversity, but not excluding ethnic diversity. Given the clear effects of name on the evaluation of CVs ^2,3,5,6^, and consistent with the League of European Research Universities (LERU) recommendations for 2-phase, partially anonymized CV evaluation,^1^ we decided to primarily focus our initial efforts on the redaction of application packages, including CVs, during faculty-level job searches. Simultaneously, we also included alterations to job advertisements to emphasize the equity, diversity, and inclusion (EDI) aspects of the application process. It is important to acknowledge that we are far from experts in this area and at the start of this process in 2020, we were largely unaware of existing literature in the area. Nevertheless, we feel that our real-world experience in this exercise could inform practices in other departments, particularly in STEM fields, since the effects of this relatively simple exercise have been overwhelmingly positive and emphasize the ability of willing units to make meaningful change in the absence of internal “expertise”, or time-expensive (and therefore access-restrictive) formal training programs.

## Methods

### Redaction and Evaluation of Application Materials

We restricted our comparative analyses to searches for Assistant Professor positions over the last 10 years, excluding the search for an external Head of Department as we were unable to obtain application pool equity data for the latter. In our department, applicants for faculty positions are requested to provide a CV, a description of their proposed research, a teaching statement, and more recently, an EDI statement. The inclusion of an EDI statement in more recent competitions was in keeping with the movement of NSERC to include better evaluation of EDI in their grant evaluation matrix. Beginning in 2020, all application packages were redacted by the Head of the Department, who, as per the Faculty Association Collective Agreement, was not directly involved in the applicant evaluation process. This was a multi-stage process. First, the applications were separated into three categories, based on our faculty association collective agreement at the time. Category 1 were candidates who were Canadian Citizens or Permanent Residents (PR), eligible (defined solely at this phase as having a PhD in any broadly related area and at least two years of post-doctoral experience), and submitting an application containing all requested materials. Category 2 applicants were not a Canadian Citizen or PR, but otherwise eligible and submitted a complete application package. The collective agreement in place during this period stipulated that category 2 candidates could only be evaluated if a qualified Canadian Citizen or PR was not identified. Category 3 candidates were either not eligible or submitted incomplete application packages. Placement in category 3 was validated by two members of the search committee after the unredacted stage of evaluation was reached.

Applications in category 1 were subjected to redaction as follows (see Figure 2 for a timeline). A duplicate of each application pdf was created and all names, all country names, all institutional names, all contact information, all leaves of absence (e.g., parental leave), the location of talks, and any information from which the candidate’s sex, gender, religion, ethnicity, race, age, nationality (beyond being either Canadian or a permanent resident), professional or personal affiliations could reasonably be discerned were permanently removed using the redaction function of Preview for Macs (Note that Adobe Acrobat Pro has a similar function). Any information (e.g., use of gendered pronouns) that could provide identifying information about trainees, co-authors, colleagues, or mentors was also redacted as these can allow for the inference of status ^17,18^. The information retained included: the location and titles of conference presentations associated with international or Canadian scientific societies, degree type and year awarded, thesis titles, and publication titles, year, and journal. Added to the CV was the location of the candidate in the authorship line (e.g., “1/5” indicates first author out of five total authors) of each publication and presentation. See Supplemental File 1 and 2 for examples of an original and redacted application.

**Figure 2.**
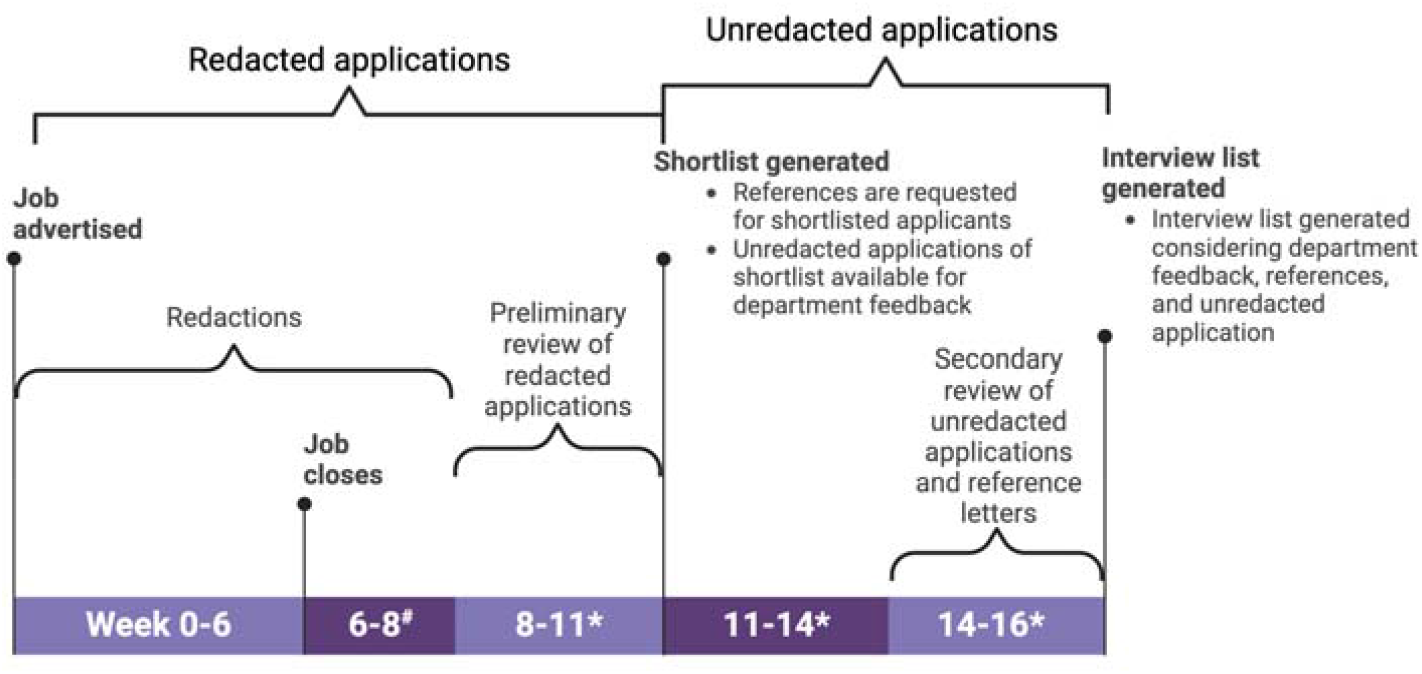
Timeline for hires following the redaction process. Redactions added approximately two weeks to the timeframe required to generate an interview list. # This is the only additional process that was added compared to without redaction. *The exact timing for these steps depends on the availability of the committee members and referees’ response speed. Created in BioRender. Christian, S. (2024) BioRender.com/h32d783

The search committee evaluated the redacted copy of each category 1 application package in order to generate a shortlist (≤ 10 candidates) for which reference letters were requested. Each member of the search committee came up with their own method of evaluation, based on the position advertisement. After this list was generated, and reference letters received, the unredacted application materials plus reference letters were provided to the search committee. Unredacted application materials, but not reference letters, were also made available to all faculty members within the department for their comments. The final interview list (typically 3-4 candidates) was then compiled by the search committee based on a re-evaluation of the initial shortlist that included the unredacted application, the letters of reference, and feedback from current faculty members.

Human Resources (HR) was able to provide aggregate equity data on all competitions based on applicants completing a self-identification checkbox questionnaire. We analyzed gender and racialized criteria as the other criteria (LGBTQ+, disabled, and Indigenous) had less than 5 applicants per search and so were repressed. Statistical analysis between the percentage of applications and interviews were done using the Student’s T-test and were considered significant at P<0.05.

### Job advertisement text

Text was added to the job advertisements to indicate the commitment of the department to EDI. For example, “The Department is committed to increasing diversity within the faculty complement and will provide additional mentorship and supports to the successful applicant as appropriate.” Or “As part of the commitment of the Department of Biochemistry to EDI, the hiring committee is implementing best practices to reduce unconscious bias and to ensure that initial shortlisting of candidates includes those from under-represented target groups at the level they exist in the applicant pool according to Statistics Canada.”

In addition, descriptions of qualifications and fields were kept as broad as possible, keeping the list of requirements to a minimum, while respecting the area of expertise required for a particular position. Moreover, we included the possibility of a spousal hire as, anecdotally, this is often a bigger consideration for women than men.

### Interview procedure

To facilitate selecting the best possible candidates a number of changes were also incorporated into our interview procedures, in an attempt to move closer toward EDI best practices. The number of meetings over the two-day interview period was reduced, to allow breaks for candidates, to further decrease stress and fatigue. Candidates were provided the opportunity to identify which individuals/groups on campus they would prefer to meet with, in addition to the Collective Agreement-mandated meetings. All interview components, including formal committee questions, were standardized within each competition. Every effort was made to present an honest and open picture of life in the Department, University, and city to candidates, with an emphasis on maintaining positive work-life balance. All candidates were provided with the opportunity to meet with a realtor as part of the interview, to provide independent information on housing and cost-of-living aspects in the area.

## Results and Discussion

### The process of redaction

The faculty debated on how best to proceed with redaction. Multiple suggestions were put forward, including requesting applicants to self-redact, potentially by establishing an online form. In the end, this was rejected as many felt that this would be an additional barrier for potential applicants. We discussed having an administrative assistant perform the redactions, as has been described previously ^19^, however, ultimately decided against this as our administrative assistants did not have the time nor expertise to judge which features and factors should be redacted or not. For example, it was agreed that redaction of the entire application, and not just the CV, was required to remove all indications for gender or country of origin. Thus, someone familiar with the academic environment (e.g., knows when an item, such as a country in the name of a journal, would likely provide identity clues) must do the redaction. This was not a trivial task as the CV, research proposal, teaching statement, and EDI statement (when requested) all needed to be redacted. In the end, the Head of the Department took on this task, as they are not allowed to be on the search committee but have the required expertise. On average, redactions took approximately 1.5 hours per application, with a range of 45 minutes to 2 hours. In no instance did post-categorization validation result in Category 3 applications being moved to either Category 1 or 2, or vice versa.

There was the occasional error in the redaction process, all of which were errors of omission (information that should have been redacted but was missed), but this occurred, on average, with at most one instance in one or two applications per competition and usually fewer than that. The search committees did not feel that this impacted their overall evaluations because, by the time they encountered the errors, they had already established their workflow for evaluation (e.g., rubrics), as discussed below, and begun forming their assessment. Furthermore, it was felt that any omissions accounted for such a small percentage of the total redacted material of individual applications, that it would only enable explicit biases to be expressed.

One concern from some faculty members was that even with rigorous redaction, applicants could be easily identified by a simple PubMed or Google Scholar search of their published articles. Therefore, the search committee members agreed that they would avoid circumventing the aim of this exercise by actively trying to identify the applicants. Only one committee member (in one search) confessed to actively looking for the identity of one or more applicants. No other committee member revealed that they actively sought out the applicants’ identities, however, there were cases where a committee member had previously supervised an applicant. In this case, the shared publication titles between the applicant and committee member made the identity of the applicant obvious to the committee member. However, none of the other committee members were able to identify the applicant. Overall, we suggest periodic reminders during the evaluation process to underline that the purpose of redaction is not to test search committee members’ ability to use search engines, but to further a goal that the department members have unanimously decided upon (i.e., to reduce bias) is important.

An unforeseen benefit to having the first round of evaluations redacted was that conflicts of interest (COI) between the search committee and the applicant pool were avoided in the first selection step. Normally, a perceived or real COI would require that the committee member recuse themselves from the entire search process. However, as the applicants were unknown to the committee as a whole, a member was able to self-recuse for the problematic application only. After this stage, the unredacted applications from the shortlist were made available to the search committee and COIs were declared at this time. Reducing the COI issue meant that applicants who did not make the shortlist did not delay search procedures while we identified replacement committee members, who may not have the same level of expertise. This is particularly important in a department such as ours where the pool of search committee members is small.

### Feedback on the Evaluation Process

Overall, search committee members felt that this process improved their ability to rate applicants. More than one person expressed that they felt relief that they no longer had to attempt to reduce their implicit bias consciously and could just focus on the qualifications. A number of committee members expressed their initial reluctance and suspicions of the process, which, they admitted were unfounded once they began their evaluations.

An unanticipated benefit was that the redaction helped reviewers focus when the candidate had completed any of their training at our institution. Prior to this, there was a prevailing trend of restricting the shortlist to no more than one “local” candidate. In addition, anecdotally, there appeared to be fewer instances where committee members would claim a “lack of fit” without substantive reasons or speculations that the candidate “may not be able to communicate well in English”. At least one initially skeptical committee member noted that while they did not feel redaction altered their assessment of applicants, it did make them more consciously aware of potential implicit biases they may have, and from that point alone it was beneficial.

The lack of identifying information necessitated the creation of evaluation rubrics by each committee member. Prior to the redaction of applications, the use of evaluation rubrics was rare. After the introduction of redacted applications, the use of rubrics became common and even expected. The use of rubrics has recently been shown to increase diversity of hiring ^20^, therefore, the self-adoption of rubrics may be one major factor in the differences seen before and after redaction. In addition, by allowing each member to create their own rubric, it maintained a diversity of opinion on the relative merits of each aspect of the application. It also improved buy-in on the importance of using a rubric at all. Thus, the redacted application has the benefit of encouraging rubric use while maintaining flexibility and diversity of opinion.

### Significantly more women were hired after the adoption of redacted applications

In order to evaluate the effects of redacted applications on hiring, we compared hiring during the periods 2010-2020 (5 hires; no redaction) and 2021-2024 (5 hires; redacted applications). We first compared the percentage of applicants to the percentage of interviews for female and racialized candidates and found no significant difference before or after redaction (Figure 3C-D). For female applications, the percent interviewed tended to follow the application pressure while for racialized applications, there was no clear pattern (Figure 3A-B). We next compared the percentage of women and individuals from racialized groups who were scored as the top candidate, and who were ultimately hired (Figure 3). We found a substantial difference in the number of women offered positions (5/5) during the redaction period compared to before the use of redaction (1/5). Similarly, the number of women ultimately hired increased to 80% (4/5) compared to 20% (1/5). The number of top-ranked racialized applicants did not substantially change with or without redaction (1/5 for both) and the number of hires was almost the same (3/5 vs 2/5). However, when the effects of retirements were combined with new hires, faculty member makeup became more diverse after the intervention. Notably, the department reached gender parity post-intervention with a trend towards increasing racial diversity, as well (Figure 1). In fact, our department is now in-line with the recommendations of the Government of Canada’s 50-30 challenge, where 50% of people in an organization are women or gender-diverse and 30% are members of other equity-deserving groups ^21^.

**Figure 3.**
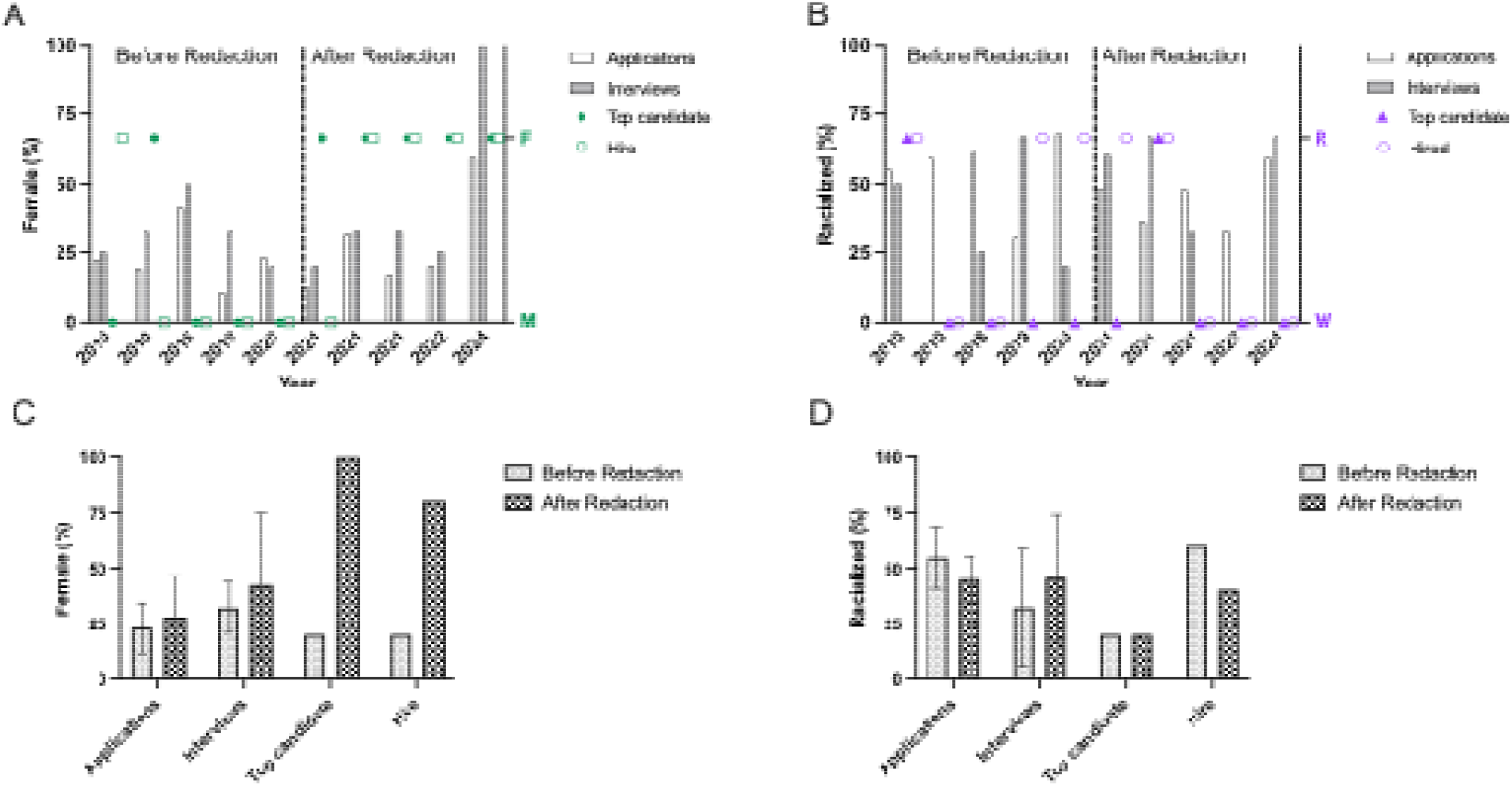
Redactions increased the number of female faculty, but not racialized faculty, hired but normalized racialized applications to be closer to the application pressure. A) Left axis: Percentage of self-identified females in each category for each hiring competition. Right axis: Identity of the top-ranked and hired candidate. B) Left axis: Percentage of self-identified racialized individuals in each category for each hiring competition. Right axis: Identity of the top-ranked and hired candidate. C) Percentage of females who applied, were interviewed, ranked the top candidate and ultimately hired. P=0.48 for applications and P=0.37 for interviews before vs. after redaction initiation. Shown are mean ± SD for applications and interviews. D) Percentage of racialized individuals who applied, were interviewed, ranked the top candidate and ultimately hired. P=0.61 for applications and P=0.54 for interviews before vs after redaction initiation. Shown are mean ± SD for applications and interviews.

It is most probable that the percentage of female candidates interviewed followed the application pressure because, under the collective agreement of the faculty association, the search committee must submit the shortlist to Human Resources (HR) for evaluation. In this submission, a justification for the exclusion/inclusion of any candidate belonging to an underrepresented group must be provided. Therefore, search committees have always been conscious of including women and racialized candidates on the interview list. Given that the end result was so few women hires, however, we believe that the women included on the interviewee list were not always seriously considered, but may have been included as “token” interviews. It is notable that the total number of applications during the non-redaction period was 41±16 while during the redaction period it was 22±6 (P=0.06 by T-test) suggesting the applicant pool was at least as deep and likely deeper during the non-redaction period. Therefore, it is very unlikely that there were fewer qualified female candidates prior to the implementation of redaction. Moreover, we believe that this suggests that the current practice where HR checks interview lists for diversity after they are compiled is insufficient to address disparities in hiring. Rather it is at the level of the hiring committee that this needs to be addressed.

With respect to racialized candidates, we found that prior to redaction, there was one search where the percentage interviewed followed the application pressure and one where exceeded the application pressure. However, after redaction, the percentage of racialized candidates who interviewed exceeded the application pressure in three out of the five searches. Therefore, there was a clear effect of redaction improving the likelihood for a racialized person to be interviewed.

One limitation of this experience is that we are unable to separate the effects of application redaction from changes to the interview processes. The percentage of applications from female and racialized candidates was very similar in the pre- and post-redaction eras so it is likely that the job advertisement text had little effect. The additional discussions around diversity within the department, because of the changes in the hiring process, may have had a direct impact, as well. It is also possible that the push to increase diversity might have had a reverse bias effect and influenced the search committee’s decisions. If this was the case, however, then we would have predicted that the quality of the hires in the redaction era would be lower than expected. Although quality is an imprecise term difficult to quantify, available evidence suggests this is not the case. All hires (3/3) from the redaction era who have applied for national tri-council (NSERC, CIHR, or SSHRC) funding for their research have received it. This compares to a 54% success rate for the “Genes, Cells, and Molecules”, “Biological Systems and Functions”, and “Chemistry” NSERC evaluation groups for early career researchers (ECR) in 2021-2024 ^22^. Moreover, two of the hires were successful in highly competitive national funding competitions with success rates of less than 15%. The fourth hire from this timeframe was in a non-traditional area (Life Sciences pedagogy), an area that remains primarily reliant on non-tri-councils funding for research program initiation and the most recent hire has not yet had an opportunity to apply for national funding. We are limited to this indicator of success at present as redaction era hires are still ECR, and it is too soon to analyze other merit indicators of success (e.g., publication record, graduate supervision success, award nominations, promotion and tenure decisions).

Overall, our findings are in line with previous reports showing that direct interventions with search committees increases the diversity of recruitment ^23–26^.

### Evaluating unredacted applications is also important

One concern that was raised early on was that the search committees are unable to perform a holistic review of applications, as has been recommended by others ^27^ and in keeping with the San Francisco Declaration on Research Assessment (DORA) recommendations ^28^. For example, one member pointed out that one applicant had significant outreach activities that had been redacted (because much of it was with women in science) and revealing the full application significantly raised their ranking. While the number of the redacted outreach activities in the CV clearly indicated that the applicant was engaged in outreach, the impact of the outreach was not fully evident until the unredacted application was assessed. Therefore, it is critical that the shortlist is evaluated a second time with fully unredacted applications before creating the interview list. In addition, the initial shortlist should be relatively long so that applicants with other activities are captured appropriately. This second look gives the search committee the ability to view the applications through a holistic lens and to take into consideration other aspects of the application.

### The question of scalability

A concern that we have heard from other departments is that this process is not scalable for searches that attract hundreds of applicants. There are a few mitigating factors that help improve feasibility with large searches. The first is the importance of eliminating ineligible or incomplete applications. In our broader experience, this removes 25-30% of all applications from consideration. If there are further guidelines for country of residence, this could further reduce the number of applications under consideration. We also recommend that applications be categorized and, when necessary, redacted as they are submitted so that the work can be accomplished in a manageable time frame (Figure 3). However, a large number of applications may require that more than one individual perform the redaction. In this case, very clear guidelines must be provided so that the redactions are consistent. Spot checks by another redactor could be performed to ensure consistency as well. Recently, Artificial Intelligence tools have become available that could also be used for this process as well. We ran the auto-redaction feature from “Redactable” to test one such program (see Supplemental File 3) and found that additional manual curation would be needed to both remove unneeded redactions and add necessary redactions. Lastly, we note that the Head of the Department performed the redaction for our hires, reinforcing the importance of having leadership buy-in and providing the appropriate recognition of the importance of the task.

## Conclusion

In conclusion, our customized intervention resulted in achieving gender parity and increased racial diversity within the department over four years. We believe that no single action was responsible for this result. Rather, it was the accumulation of increased awareness and active intervention that combined to increase the diversity within the Department.

## Acknowledgements

We would like to thank Mandy Penney and Amanda Brace from Human Resources at Memorial University of Newfoundland for their help in retrieving and aggregating the equity data on the applicants. We thank NSERC for their continued support of EDI initiatives by tracking equity data in their award competitions. We are grateful for the enthusiasm and participation of the entire Department of Biochemistry but in particular Drs. Janet Brunton, Katie Wilson and Robert Bertolo for constructive feedback on initial drafts of this submission. Graphical abstract created in BioRender. Christian, S. (2024) BioRender.com/k60p368.

