## Supplementary material for "Equity in Action: A Four-Year journey towards Gender Parity and Racial Diversity in Biochemistry Hiring": Figure S1

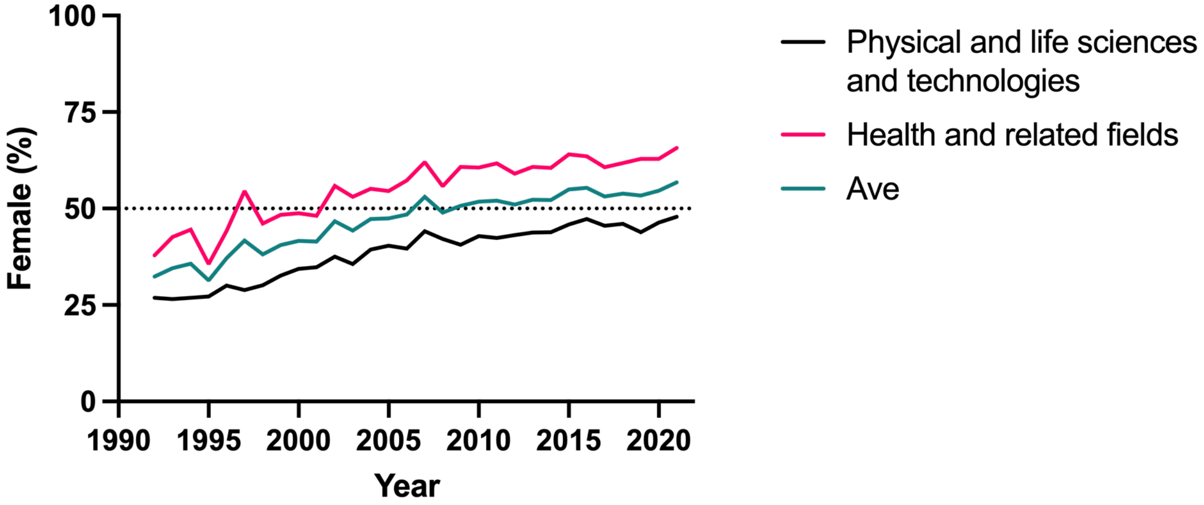


Figure S1. Percentage of females earning a Doctoral Degree or equivalent in Canada from 1992 to 2021 (Statistics Canada).
Table 37-10-0135-02 Proportion of male and female postsecondary graduates, by field of study and International Standard Classification of Education. https://www150.statcan.gc.ca/t1/tbl1/en/tv.action?pid=3710013502 (2021).
