## Supplemental File 1 and 2 for "Equity in Action: A Four-Year journey towards Gender Parity and Racial Diversity in Biochemistry Hiring"

### CURRICULUM VITAE

[redacted]

Name: [redacted]

[redacted]

Citizenship: [redacted]

[redacted]

#### Contact Information:

[redacted]

#### Current Position:

Postdoctoral Research Fellow

Department of Nutritional Sciences, [redacted]

2019 - Present

#### Education:

Ph.D. in Nutritional Sciences

[redacted] 2019

Dissertation: "Nutrient-Gene Interactions in Metabolic Regulation"

**Scholarship:** [redacted] Research Fellowship

M.Sc. in Molecular Biology

[redacted] 2014

Thesis: "Epigenetic Control of Metabolic Pathways"

B.Sc. in Biochemistry

[redacted] 2012

**Scholarships:** [redacted] Scholarship, [redacted] Alumni Entrance Scholarship, Undergraduate Research Fellowship

#### Research Interests:

My research focuses on the complex interactions between nutrients and genes, particularly in the context of metabolic regulation and chronic disease prevention. I am particularly interested in how nutrient sensing pathways influence gene expression and how these interactions can be leveraged to develop personalized nutrition strategies. My work has explored the epigenetic regulation of metabolic pathways and the role of specific dietary components in modulating these processes. I have been recognized for my contributions to this field, having received multiple awards from the prestigious [redacted] granting organization for studies that have significantly advanced our understanding of nutrient-gene interactions. These accolades have underscored the impact of my research and provided essential support for the development of innovative approaches to studying these critical biological mechanisms and training of Highly Qualified Personnel from diverse backgrounds.

**Research Interest Summary:**

- Nutrient and gene interactions
- Metabolic pathways and nutrient sensing
- Epigenetic regulation of gene expression
- Nutrient-gene interactions in chronic diseases

#### Publications:

1=/3+ 1. [redacted] et al., "Nutrient-Gene Interaction in Liver Metabolism," Journal of Nutrient-Gene Interactions, vol. 26, no. 6, pp. 110-119, 2022. (*Contributed equally)

1/3 2. [redacted], "Regulation of Metabolic Pathways by Dietary Components indigenous to [redacted]" Metabolism and Gene Regulation, vol. 56, no. 10, pp. 662-671, 2021.

1/6 3. [redacted], "Nutrient Sensing Mechanisms in Cells," Journal of Nutrigenomics, vol. 31, no. 2, pp. 99-106, 2020.

5/5 4. [redacted], "Epigenetic Changes Induced by Nutrients," Genes and Nutrition, vol. 20, no. 10, pp. 2102-2112, 2019.

5. [redacted]. "Dietary Influence
4/5 on Gene Expression in [redacted] high performance athletes," [redacted] Journal of Molecular Nutrition & Food Research, vol. 27, no. 3, pp. 81-84, 2018.

#### Conference Presentations:

1. "Nutrient Sensing and Gene Regulation," Nutrigenomics Conference, [redacted], 2023.
2. "Diet and Epigenetics," Metabolism Conference, [redacted] 2022.
3. "Nutrient-Gene Interactions," Genomics Symposium, [redacted], 2021.
4. "Metabolic Pathways Regulation," [redacted] Biology Conference, [redacted], 2020.
5. "Dietary Impacts on Gene Expression," [redacted] Metabolic Science Meeting, [redacted] 2019.

#### Equity, Diversity, and Inclusion Statement:

As a researcher and educator, I am committed to fostering an inclusive and diverse environment within my laboratory and beyond. I believe that diversity in perspectives, experiences, and backgrounds is essential for driving innovation and excellence in scientific research. To this end, I actively strive to recruit and support a diverse group of students and researchers in my lab, ensuring that individuals from all walks of life have the opportunity to contribute to and benefit from our collective work.

In my research, I make it a priority to account for sex differences in study designs, recognizing the critical importance of understanding how biological factors can influence health outcomes across different populations. This approach not only enhances the rigor and relevance of our findings but also ensures that our research addresses the needs of diverse communities.

To further my commitment to equity and inclusion, I have completed unconscious bias training, which has equipped me with the tools to recognize and mitigate biases in my professional interactions and decision-making processes. Additionally, I volunteer with the [redacted] Child Refugee Education Program, an organization dedicated to helping children from lower socioeconomic backgrounds access STEM enrichment programs. Through this work, I aim to inspire and support the next generation of scientists, particularly those who may face barriers to pursuing careers in STEM fields. [redacted]

[redacted]. By providing mentorship and resources, I hope to help these students succeed in high school and university, ultimately contributing to a more diverse and inclusive scientific community.

#### Research Experience:

**Postdoctoral Research Fellow**

[redacted]

2019 - Present

- Conducted advanced research on nutrient and gene interactions
- Developed and implemented experimental protocols to investigate metabolic pathways
- Published findings in high-impact peer-reviewed journals
- Mentored graduate and undergraduate students in the laboratory
- **Research Grant:** [redacted] Health Research Grant

**Graduate Research Assistant**

[redacted]

2014 - 2019

- Conducted research on epigenetic regulation of gene expression
- Assisted in grant writing and securing funding for research projects
- Collaborated with interdisciplinary teams to advance research goals

#### Teaching Experience:

Instructor, Advanced Nutrigenomics

[redacted]

Fall 2022

- Designed and delivered lectures on nutrient-gene interactions
- Created and graded assessments to evaluate student learning

Teaching Assistant, Molecular Biology

[redacted]

2015 - 2018

- Assisted in course instruction and lab supervision

- Provided support and feedback to students on assignments and projects

#### Professional Affiliations:

- Member, [redacted] Nutritional Sciences Society
- Member, [redacted] Molecular Biology Association
- [redacted] Society for Nutrition

#### Awards and Honors:

- [redacted] Research Award, 2022
- [redacted] Research Fellowship, 2019
- [redacted] Scholarship, 2012
- Undergraduate Research Fellowship, 2011

#### Students Supervised:

- 2024 [redacted] ([redacted] Research Experience Program)
- 2023 D. Scully ([redacted] Society for Nutrition "Nutrition: The Truth is Out There" Award)
- 2022 [redacted] ([redacted] Undergraduate Career Research Experience Program)
- 2020 [redacted] ([redacted] Natural Sciences Undergraduate Research Award)

#### References:

Available upon request.
