## Supplemental File 3 for "Equity in Action: A Four-Year journey towards Gender Parity and Racial Diversity in Biochemistry Hiring"

### CURRICULUM VITAE

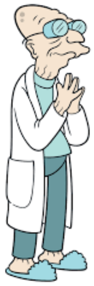

Name: [redacted]

Pronouns: He/Him

Marital Status: Separated

Citizenship: Martian Permanent resident, [redacted]

Date of Birth: [redacted]

#### Contact Information:

[redacted]

#### Current Position:

[redacted] Research [redacted]

[redacted]

[redacted] - Present

#### Education:

Ph.D. in [redacted]

[redacted], [redacted]

Dissertation: "Nutrient-Gene Interactions in Metabolic Regulation"

**Scholarship:** Interplanetary Research Fellowship

M.Sc. in Molecular Biology

[redacted], [redacted]

Thesis: "Epigenetic Control of Metabolic Pathways"

B.Sc. in Biochemistry

[redacted], [redacted]

**Scholarships:** Fraternity of Galactic Scientists Scholarship, Springfield All-Boys High School Alumni Entrance Scholarship, Undergraduate Research Fellowship

#### Research Interests:

My research focuses on the complex interactions between nutrients and genes, particularly in the context of metabolic regulation and chronic disease prevention. I am particularly interested in how nutrient sensing pathways influence gene expression and how these interactions can be leveraged to develop personalized nutrition strategies. My work has explored the epigenetic regulation of metabolic pathways and the role of specific dietary components in modulating these processes. I have been recognized for my contributions to this field, having received multiple awards from the prestigious [redacted] granting organization for studies that have significantly advanced our understanding of nutrient-gene interactions. These accolades have underscored the impact of my research and provided essential support for the development of innovative approaches to studying these critical biological mechanisms and training of Highly Qualified Personnel from diverse backgrounds.

**Research Interest Summary:**

- Nutrient and gene interactions
- Metabolic pathways and nutrient sensing
- Epigenetic regulation of gene expression
- Nutrient-gene interactions in chronic diseases

#### Publications:

1. [redacted], [redacted]*, [redacted] et al., "Nutrient-Gene Interaction in Liver Metabolism," Journal of Nutrient-Gene Interactions, vol. 26, [redacted], pp. [redacted], [redacted]. (*Contributed equally)

2. [redacted], [redacted], and [redacted], "Regulation of Metabolic Pathways by Dietary Components indigenous to [redacted] & Labrador," Metabolism and Gene Regulation, vol[redacted] [redacted], no. 10, pp. 662-671, [redacted].
3. [redacted], [redacted], [redacted], [redacted] and [redacted] [redacted], "Nutrient Sensing Mechanisms in Cells," Journal of Nutrigenomics, vol[redacted], [redacted], pp. 99-106, [redacted].
4. [redacted], [redacted], [redacted] and [redacted], "Epigenetic Changes Induced by Nutrients," Genes and Nutrition, [redacted], [redacted], pp. 2102-2112, [redacted].
5. [redacted], [redacted], [redacted], and [redacted] "Dietary Influence on Gene Expression in Australian female high performance athletes," Australian [redacted] [redacted] [redacted], [redacted], pp. 81-84, [redacted].

#### Conference Presentations:

[redacted] "Nutrient Sensing and Gene Regulation," Nutrigenomics Conference, New York, [redacted]
2. "Diet and Epigenetics," Metabolism Conference, [redacted] ity, [redacted]
3. "Nutrient-Gene Interactions," Genomics Symposium, Jupiter Station, [redacted]
[redacted]. "Metabolic Pathways Regulation," Interplanetary Biology Conference, Saturn Ring, [redacted]
5. "Dietary Impacts on Gene Expression," Venus Metabolic Science Meeting, Venusville, [redacted]

**Graduate Research Assistant**

[redacted]

[redacted] - [redacted]

- Conducted research on epigenetic regulation of gene expression
- Assisted in grant writing and securing funding for research projects
- Collaborated with interdisciplinary teams to advance research goals

#### Teaching Experience:

Instructor, Advanced Nutrigenomics

[redacted]

[redacted]

- Designed and delivered lectures on nutrient-gene interactions
- Created and graded assessments to evaluate student learning

Teaching Assistant, Molecular Biology

[redacted]

[redacted] - [redacted]

- Assisted in course instruction and lab supervision

- Provided support and feedback to students on assignments and projects

#### Professional Affiliations:

- Member, [redacted]
- Member, [redacted]
- [redacted]

#### Awards and Honors:

- [redacted] Research Award, [redacted]
- Interplanetary Research Fellowship, [redacted]
- Fraternity of Galactic Scientists Scholarship, [redacted]
- Undergraduate Research Fellowship, [redacted]

#### Students Supervised:

- [redacted] (STEM for Girls Research Experience Program)
- [redacted] ([redacted] "Nutrition: The Truth is Out There" Award)
- [redacted] and [redacted] (Futurama [redacted] Undergraduate Career Research Experience Program)
- [redacted] (Martian Natural Sciences Undergraduate Research Award)

#### References:

Available upon request.
